# StomaVision: stomatal trait analysis through deep learning

**DOI:** 10.1101/2024.04.24.590919

**Authors:** Ting-Li Wu, Po-Yu Chen, Xiaofei Du, Heiru Wu, Jheng-Yang Ou, Po-Xing Zheng, Yu-Lin Wu, Ruei-Shiuan Wang, Te-Chang Hsu, Chen-Yu Lin, Wei-Yang Lin, Ping-Lin Chang, Chin-Min Kimmy Ho, Yao-Cheng Lin

## Abstract

StomaVision is an automated tool designed for high-throughput detection and measurement of stomatal traits, such as stomatal number, pore size, and closure rate. It provides insights into plant responses to environmental cues, streamlining the analysis of micrographs from field-grown plants across various species, including monocots and dicots. Enhanced by a novel collection method that utilizes video recording, StomaVision increases the number of captured images for robust statistical analysis. Accessible via an intuitive web interface at <https://stomavision.streamlit.app/> and available for local use in a containerized environment at <https://github.com/YaoChengLab/StomaVision>, this tool ensures long-term usability by minimizing the impact of software updates and maintaining functionality with minimal setup requirements. The application of StomaVision has provided significant physiological insights, such as variations in stomatal density, opening rates, and total pore area under heat stress. These traits correlate with critical physiological processes, including gas exchange, carbon assimilation, and water use efficiency, demonstrating the tool’s utility in advancing our understanding of plant physiology. The ability of StomaVision to identify differences in responses to varying durations of heat treatment highlights its value in plant science research.

**Plain language summary:** StomaVision is a tool that automatically counts and measures tiny openings on plant leaves, helping us learn how plants deal with their surroundings. It is easy to use and works well with various plant species. This tool helps scientists see how plants change under stress, making plant research easier and more accurate.

## Introduction

Stomata are microscopic pores on most plant aerial parts that first appeared over 400 million years ago, enabling plants to colonize terrestrial environments (Clark *et al*., 2022). Except for submerged aquatic angiosperms, plants use stomata to regulate the transpiration rate of water loss and diffusion of CO_2_ into the leaves used in photosynthesis (Olsen *et al*., 2016; Clark *et al*., 2022). Stomata open to facilitate leaf cooling through water evaporation at high temperatures and close at night or during drought to minimize water loss. The opening and closing of stomata are strongly regulated by air temperature and humidity through specialized guard cells. Stomatal anatomy, such as shape, density, area, and distribution of stomata between the abaxial and adaxial regions, are controlled by intrinsic genetic variations and extrinsic environmental factors such as temperature, photoperiod, humidity, and soil water potential (Roelfsema & Hedrich, 2005; Shimazaki *et al*., 2007; Li *et al*., 2013; Lawson & Matthews, 2020). The regulation of stomata is crucial for both plant growth and global cycles of water, energy, and carbon (Lawson & Matthews, 2020). The function and structure of stomata play a crucial role in determining stomatal conductance (*g_sw_*), assimilation rate (*A*), and water use efficiency (WUE).

Stomatal conductance (*g_sw_*) refers to the maximum rate at which gases can diffuse through the stomata, and is influenced by stomatal density and pore size. This value serves as an indicator of a plant’s ability to exchange gases through its leaves. Stomata typically cover 0.3-5% of the leaf surface area and facilitate approximately 95% of gas exchange through these pores (Kane *et al*., 2020; Lawson & Matthews, 2020). Stomatal density and patterning are both under environmental and genetic control (Serna & Fenoll, 1997; Sugano *et al*., 2010; Xie *et al*., 2022). Stomatal opening is stimulated by light and the accumulation of potassium ions (K^+^), which is mediated by electrogenic proton pumps in the plasma membrane of guard cells (Shimazaki *et al*., 2007). Guard cells also respond to changes in air humidity and the water status of distant tissues by signaling molecules such as phytohormones and hydrogen peroxide (Roelfsema & Hedrich, 2005). Enhancing our understanding of stomatal behavior and its efficiency in CO_2_ and water use is a key goal in improving crop resilience in changing climates (Bailey-Serres *et al*., 2019).

Various methods have been developed to study stomata because of their importance in plant biology; however, they remain laborious and time-consuming. Non-destructive methods, such as infrared gas exchange analyzers (Garen *et al*., 2022) and infrared imaging (Jones *et al*., 2002), infer stomatal activities through infrared gas sensors or leaf surface temperature. Infrared gas exchange analyzers are often considered the ‘gold standard’ for determining gaseous fluxes (Driever *et al*., 2023). Destructive techniques, such as stomatal imprints (Scarpeci *et al*., 2017), chemical dye staining (Eisele *et al*., 2016), and scanning electron microscopy (Tsai *et al*., 2022), provide direct visualization of stomatal traits such as stomatal density, stomatal shape, stomatal pore size, and stomatal distribution. Non-destructive and destructive methods provide complementary aspects of stomatal traits and the corresponding plant physiological status.

The choice of techniques and instruments used to measure stomatal traits was influenced by experimental design restrictions. Portable gas exchange analyzers offer a non-destructive method for continuously monitoring physiological changes over time (Durand *et al*., 2022; Wall *et al*., 2022; Xie *et al*., 2022). These instruments can measure multiple plant physiological traits, such as leaf temperature, stomatal conductance (*g_s_*), transpiration (*E*), water vapor deficit (VPD), and CO_2_ assimilation rates (A), which are critical for understanding plant physiological responses to the environment. However, these instruments are often costly and can only be used to monitor one plant at a time. Therefore, stomatal imaging is currently the most commonly used large-scale approach, particularly in the field environment, to rapidly preserve stomatal behavior before the stomatal aperture is adjusted to the environment (Lawson & Blatt, 2014; Sakoda *et al*., 2019; Bheemanahalli *et al*., 2021). However, stomatal morphology varies in terms of the size, shape, and arrangement on the leaf surface. In monocots, stomata are usually aligned in rows along veins, whereas in dicots, they can be either dispersed or clustered. Guard cells have two primary shapes, gramineous (dumbbell-shaped) and kidney-shaped, and are often accompanied by subsidiary cells (Lawson & Matthews, 2020). Traits such as stomatal size, density, and closure rate serve as effective indicators for assessing how plants respond to environmental stress and quantifying stomatal gas and water exchange parameters (Xie *et al*., 2022). Furthermore, the analysis of a large number of stomatal images is necessary for statistical robustness, and this process is both time-consuming and prone to human error (Eisele *et al*., 2016; Bheemanahalli *et al*., 2021; Xie *et al*., 2021).

Machine-aided methods to detect stomata have been developed along with the advancement of computer vision technology and improvements in imaging sensors since the 1980s (Omasa & Onoe, 1984). Classical rule-based methods rely on a large number of high-quality, manually labeled image datasets to extract specific leaf and stomatal traits using predefined thresholds. To alleviate the rigid assumptions made by rule-based approaches, machine-learning (ML)-based methods have been developed to meet the special needs of stomatal image analysis (Kutsuna *et al*., 2012; Laga *et al*., 2014; Vialet-Chabrand & Brendel, 2014; Duarte *et al*., 2017; Jayakody *et al*., 2017; Bourdais *et al*., 2019). These approaches exploit Convolutional Neural Networks (CNN), one of the most important computer vision foundation techniques, to learn and extract stomatal features from images. However, these methods are often limited to the specific microscopic settings and plant species used to train the algorithm. Recent advancements in Deep Neural Networks (DNNs) have significantly improved the performance of classic computer vision tasks for image classification, object detection, semantic segmentation, and instance segmentation in plant phenotype analysis (Miao *et al*., 2021; Tu *et al*., 2022; Wang *et al*., 2022). The use of DNNs to identify stomata has shown improved performance in various tasks (Toda *et al*., 2018; Fetter *et al*., 2019; Li *et al*., 2019, 2022; Sakoda *et al*., 2019; Aono *et al*., 2021; Jayakody *et al*., 2021; Zhu *et al*., 2021; Liang *et al*., 2022; Pathoumthong *et al*., 2023; Sai *et al*., 2023). Nevertheless, there are still significant limitations that impede their adoption and application. One major limitation is that most published methods can only detect (count) stomata (Fetter *et al*., 2019; Sakoda *et al*., 2019; Aono *et al*., 2021; Jayakody *et al*., 2021; Zhu *et al*., 2021) and have a limited ability to quantify critical stomatal parameters, such as aspect ratio, pore size, and dynamic states (open versus closed) (Jayakody *et al*., 2017; Toda *et al*., 2018; Bheemanahalli *et al*., 2021; Liang *et al*., 2022; Li *et al*., 2022; Pathoumthong *et al*., 2023; Sai *et al*., 2023; Wang *et al*., 2024). This limited functionality restricts the depth of physiological insights gained from these analyses. Many of these methods are tailored to specific species and microscopic setups, which can result in compromised performance when applied to different species or under various imaging conditions. Additionally, many of these tools do not provide versatile and user-friendly interfaces for a broader biological community (Table 1).

**Table 1.**
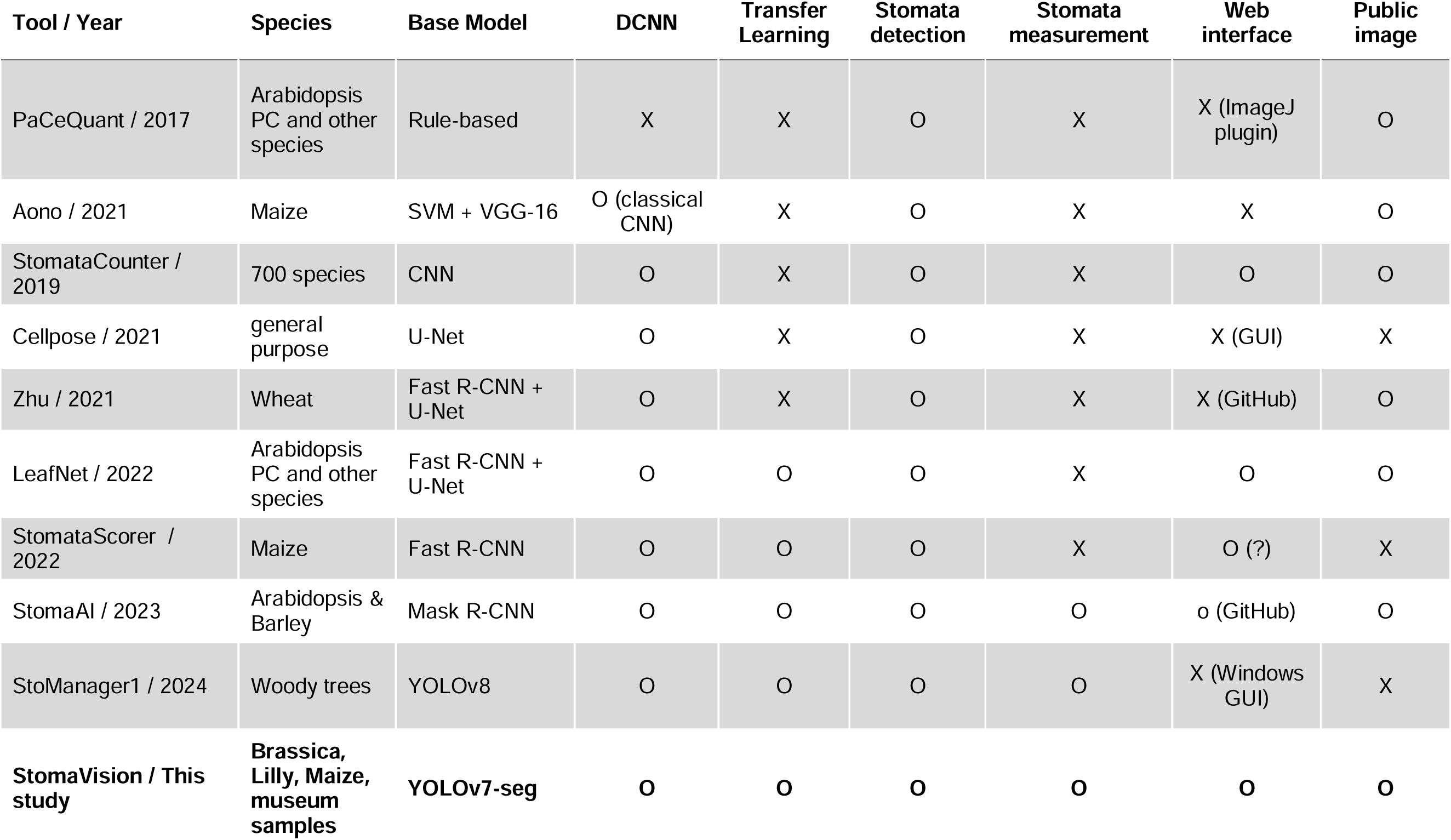
Recent research on stomata detection and measurement using machine learning.

In this study, we present StomaVision, a new online stomata detection and measurement tool that is highly accurate, robust, automatic, and high-throughput. It enables stomatal counting and pore measurement using microscope images or videos for a wide range of plant species by integrating an accurate real-time object detection model for computer vision YOLOv7 (Wang *et al*., 2023a) and a versatile data pipeline (VDP, https://github.com/instill-ai/vdp) to seamlessly connect machine learning models. To identify the bounding box of the stomata, measure the length of the two axes, and measure the pore area of stomata pores, we fine-tuned the instance segmentation model (i.e., YOLOv7-seg) using a dataset of stomata images from cauliflower, lily, maize, and images collected in the botanical garden with a wide range of species. Our evaluation results demonstrate that StomaVision outperforms models based on Mask R-CNN (He *et al*., 2017; Wu *et al*., 2020), achieving higher box average precision (AP) and mask AP accuracies. Taking advantage of the light weight and high performance of YOLOv7, we further show that using stomata micrographs with high-resolution videos can significantly reduce the time required for micrograph acquisition.

Finally, we developed an easy-to-access web-based user interface for online stomatal trait analysis and hosted it on an open-source Python platform (Streamlit, https://stomavision.streamlit.app/). Open-source code and well-documented manuals enable experienced users to install pipelines locally. Using StomaVision, we compared the stomatal traits of cauliflowers under prolonged heat stress conditions. We combined stomatal imaging and gas exchange analyzers to study stomatal and plant physiological traits, and showed a strong association between these complementary analyses. StomaVision provides an easy-to-use and affordable alternative for large-scale stomatal studies and offers a valuable resource for researchers in this field.

## Materials and methods

### Plant materials and data collection methods

*Brassica oleracea* var. *botrytis* (cauliflower) plants with diverse genetic backgrounds were cultivated in the field with average temperatures exceeding 32°C during the growth period. Fresh epidermal peels were used as plant materials for image capture. The abaxial leaf surface was immediately coated in the field with nail polish upon detachment from the stem following the methodology adapted from Scarpeci *et al*. (Scarpeci *et al*., 2017). The samples were stored at room temperature for subsequent analysis. Imprint micrographs were acquired using two light microscopes. A Carl Zeiss Axiocam 105 color digital camera with an Axiover 40 CFL inverted light microscope (Carl Zeiss) with 20x objective lens and a Carl Zeiss Axiocam 506 color digital camera with an Axio Scope A1 upright light microscope (Carl Zeiss) with 20x objective lens. A minimum of three images were captured from different locations on each imprint with resolutions of 2560 × 1920 pixels for Axiover 40 and 2752 × 2208 pixels for Axio Scope A1. To ensure representative sampling, the images were randomly cropped into smaller regions measuring 1200 × 900 pixels and 1300 × 1100 pixels (Fig. 1). To broaden the scope of the studied species, we incorporated three published datasets: Lilly (Wu, 2016), Maize (Aono *et al*., 2021), and images used by Li *et al*. (Li *et al*., 2022) and is referred to as ‘LeafNet’. It is worth noting that part of the LeafNet dataset was initially compiled to train StomataCounter (Fetter *et al*., 2019), which includes the Cuticle Database (https://cuticledb.eesi.psu.edu/) (Barclay *et al*., 2007), a Ginkgo common garden experiment (Barclay & Wing, 2016), and material from the Smithsonian National Museum of Natural History (USNM) and the United States Botanic Garden (USBG). The initial collection contained more than 6000 images of both the abaxial and adaxial cuticles in a single image (https://doi.org/10.5061/dryad.kh2gv5f).

**Fig. 1.**
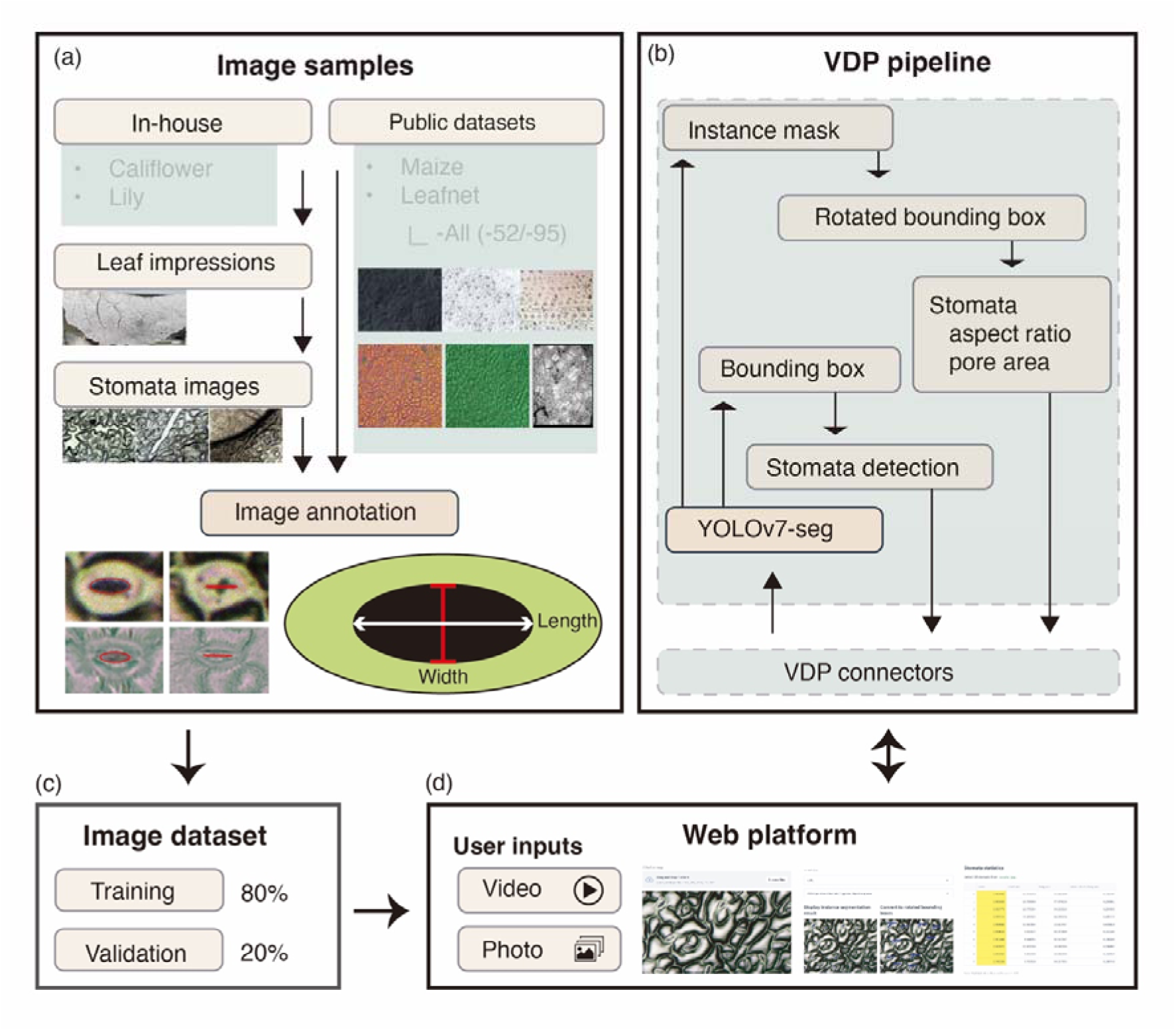
StomaVision training and inference workflow. (a) Image acquisition and stomatal annotation: Sample images were collected from independent sources to represent the different species. Graphical illustration indicating the manually labeled stomatal axes using red and white arrows, and delineating the stomatal pore with a black mask. (b) Stomatal image inference: A versatile data pipeline (VDP) was utilized to streamline the integration of image processing and machine learning algorithms. (c) Annotated stoma images were allocated to training (80%) and validation (20%) datasets. (d) User Interface: A web-based platform capable of batch processing of various file types, including MP4 videos and PNG/JPG images. It presents a side-by-side view of the original and predicted images, along with a detailed output table enumerating stomatal parameters, such as ID, image dimensions, axis lengths, axis ratio, and pore area.

For video recording, continuous imprint micrographs were acquired on a Nikon ECLIPSE Ti2 inverted microscope using a 20X objective lens and a color digital camera with a resolution of 1920 × 1080 pixels. The XY-axis of the stage and focus were manually adjusted according to the field of view during the recording.

### Data annotation and image datasets

Following the random cropping of smaller regions from the original micrograph images, the dimensions of the stomatal pores, specifically their length and width, were manually measured using ImageJ software (Schneider *et al*., 2012) to establish a ground truth dataset (Fig. 1). To facilitate the transfer of this ground truth data into a machine learning model, we used the Semantic Segmentation with EllipseLabels configuration in LabelStudio (V1.5.0) ((Tkachenko *et al*.) https://labelstud.io/, last accessed Feb 3, 2024).

Images for annotation were randomly selected from four datasets, resulting in 310 annotated images. The stomatal annotation contained various aperture features such as pore long and short axis length, pore area, and stomatal count (SOM Table 1). Additionally, we calculated the width-to-length ratio of each stoma to categorize each as either “open” or “closed,” based on a self-defined threshold. All annotations from LabelStudio were exported in the JSON format for subsequent analyses and model training.

To assess the performance of StomaVision, four distinct datasets were used. Each dataset was partitioned into training and validation subsets in an 8:2 ratio. The ‘Cauliflower’ dataset comprised 102 images, with 81 allocated for model fine-tuning and 21 for validation. The ‘Maize’ dataset contained 30 images, of which 24 were designated for model fine-tuning and six for validation (Aono *et al*., 2021). The ‘Lily’ dataset included 31 images, with 24 used for model fine-tuning and seven for validation (Wu, 2016). Finally, the LeafNet dataset (Li *et al*., 2022) consisted of 147 images, 118 of which were used for model fine-tuning and 29 for validation (referred to as ‘LeafNet-All’). These 147 images were further divided into two smaller datasets, ‘LeafNet-95’ and ‘LeafNet-52’, containing 95 and 52 images, respectively. We ensured consistency in the training and validation sets by maintaining the validation subsets in both ‘LeafNet-95’ and ‘LeafNet-52’, which were derived from the original LeafNet-All validation sets. These images were randomly selected, but owing to the format incompatibility between the data labeling methods, we manually re-labeled these images.

### StomaVision machine learning pipeline and deployment

We assessed the applicability of two leading models, Detectron2 (Wu *et al*., 2020) and YOLOv7 (Wang *et al*., 2023a), for integration into StomaVision. Detectron2 is an open-source framework developed by Facebook’s Artificial Intelligence Research Division (FAIR), employs a Mask R-CNN (He *et al*., 2017) model, and provides a comprehensive ‘Model Zoo’ for additional fine-tuning (https://github.com/facebookresearch/detectron2, last accessed Nov 3, 2023). YOLOv7, on the other hand, is versatile in handling various computer vision tasks including object detection, keypoint detection, and instance segmentation. It uses an extensive number of data augmentation techniques (e.g., scale, translation, mosaic, and mixup) during its training and fine-tuning processes. Users can fine-tune up to 28 hyperparameters to optimize the model performance. To broaden the diversity of our training dataset, we evaluated three distinct data augmentation profiles developed using Ultralytics (https://github.com/ultralytics/ultralytics). These profiles were identified based on the degree of augmentation applied, with profiles containing high, medium (referred to as “med”), and low levels of augmentation that have been extensively tested across a variety of scenarios. After conducting hyperparameter tuning over more than 32,000 iterations, we fine-tuned YOLOv7-seg with the “med” data-augmentation strategy as the foundational model for StomaVision.

The primary objective of this study was to identify stomata, compute their aspect ratios, and measure the pore area. We formulated this as an instance segmentation problem, which is a specialized form of image segmentation that focuses on detecting individual object instances and delineating their boundaries (Minaee *et al*., 2020). YOLOv7 has been shown to be effective in various computer vision applications and particularly useful for identifying small objects (Wang *et al*., 2023a). The machine learning pipeline in StomaVision comprises two main modules. The first is an instance segmentation module that identifies stomata (Fig. 1). The segmentation masks generated by this module were subsequently converted into rotated bounding boxes to calculate the aspect ratio of each detected stoma. The model was fine-tuned on an NVIDIA DGX workstation (NVIDIA) equipped with a 64-core CPU, 192 GB of memory, and four NVIDIA V100 GPUs. YOLOv7 was fine-tuned across multiple datasets, including the ‘Maize’, ‘Lily’, ‘LeafNet’, and ‘Cauliflower’ datasets, to evaluate its few-shot learning capability (Xian *et al*., 2017; Song *et al*., 2022). We set the maximum epoch to 300 and implemented early stopping to mitigate overfitting and to reduce the likelihood of model collapse.

To demonstrate StomaVision’s versatility and ease of access for plant biologists, we integrated it into a Versatile Data Pipeline (VDP) (Fig. 1). VDP is an open-source tool that enables the deployment of robust data and machine learning pipelines with minimal overhead. The StomaVision website, constructed using Streamlit (https://github.com/streamlit/streamlit), was deployed on the VDP cloud platform, allowing users to directly access only demos using their own stomata images. To further ensure long-term sustainability, StomaVision was packaged in a Docker container, with detailed setup instructions for local instances and training guidelines for species-specific model development.

### Evaluation metrics

To evaluate the performance of StomaVision, we used multiple evaluation metrics to assess the prediction performance. We incorporated a general approach by comparing the number of stomata identified in humans and machines (Fetter *et al*., 2019; Sai *et al*., 2023). The prediction of stomata pixels could have four potential results: true positive (TP) represents the number of pixels that have been correctly classified as stomata; false positive (FP) represents the number of background pixels misclassified as stomata; false negative (FN) represents the number of pixels misclassified as background; and true negative (TN) represents the number of background pixels that have been properly classified as background.

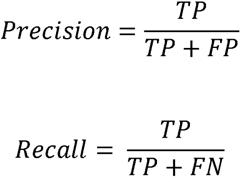

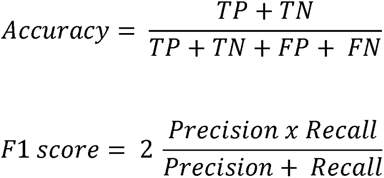

Because StomaVision can predict the stomata area to the image pixel level, we used a more sophisticated method to evaluate the prediction and ground truth data. The average precision (AP) metrics were defined in the MS-COCO (Microsoft Common Objects in Context) Challenge and are widely accepted by the computer vision community to evaluate object detection performance (https://cocodataset.org/#detection-eval) (Lin *et al*., 2014). Average precision refers to the weighted mean of the precision achieved at each threshold, which combines recall and precision to give a single score for a model’s ability to correctly identify positive instances.

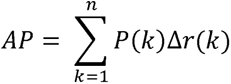

Where P(k) is the precision at the *k*^th^ threshold, and Δr(k) is the change in recall from the (k-1)^th^ to the *k^th^* threshold.

The bounding box average precision (box AP) and mask average precision (mask AP) represent the average detection precision using bounding boxes and masks under different intersections over union (IoU) thresholds. The IoU score is also known as the Jaccard Index.

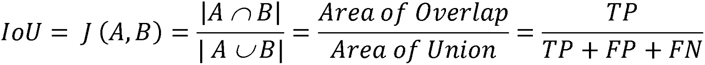

AP50 represents AP at an IoU threshold of 0.5 (50% of the area) overlap.

### Plant growth, heat stress treatment and plant physiology measurements

An elite F1 hybrid line *Brassica oleracea* var. *botrytis* cv. H37 (Ching Long Seed Co. Ltd., Taiwan) was used to examine the effects of heat stress on plant growth and physiology. The seedlings were grown in soil in a climate-controlled chamber with a 16-h day length provided by fluorescent light at a photosynthetic photon flux density (PPFD) of 360 ± 75 µmol·m^−2^·d^−1^ and day/night temperatures of 26/23°C. For heat treatment, 17-day-old plants were transferred to a growth chamber with a 16-h day length provided by LED light at 360 ± 75 µmol·m^−2^·d^−1^ and day/night temperatures of 40/23°C for up to 10 days. To better understand the impact of different heat stress durations, heat treatment was divided into short-term (2-4 day) and long-term (6-10 day) groups. After the heat treatment, the plants were transferred back to the original climate-controlled chamber with day/night temperatures of 26/23°C to record the curd induction time.

To assess the effects of heat stress on stomatal conductance and leaf physiological traits, stomatal micrographs, plant phenotypes, and physiological traits were assessed every two days before and during the heat treatment on fully expanded mature leaves during Zeitgeber time 5-7. Stomatal images were collected from the abaxial leaf surface and fixed using a nail polisher as described above. Stomatal micrographs were analyzed using the StomaVision pipeline. Thermal images were captured using a FLIR E8 infrared camera (Teledyne FLIR LLC). Stomatal conductance and leaf physiological traits were measured using a gas exchange analyzer (LI-COR 6800, LI-COR Inc.) under chamber conditions set to a CO_2_ concentration of 400 μmol/mol, light intensity of 1,000 µmol·m^−2^·d^−1^, and relative humidity of 70%, with steady-state waiting times of 5–10 min. The chamber temperature was set according to the experimental group, with the control group set to 26°C and the heat-treatment group set to 40°C. Intrinsic water use efficiency (iWUE) was calculated as the ratio of the net rate of CO_2_ assimilation (A) to stomatal conductance (*gsw*). Plant physiological traits involved in enery exchange, gas exchange and photosynthsis are listed in SOM Table 1 (Xie *et al*., 2022).

## Results

### Stomata micrographs

The inherent limitations of machine learning techniques require high-quality, extensive, and diverse training data to ensure model generalization across different image types. Our training dataset primarily consisted of cauliflower plants with a total of 102 cauliflower images, which feature a classical dicot stomatal structure, were collected under field conditions in a time-constrained and limited operational space. The stomatal micrographs, derived from a realistic real-world setting, may have introduced increased noise during model training and prediction, potentially reducing accuracy. However, this approach creates a scenario that accurately reflects the general applicability of the program in practical contexts.

To create a generalized model that can process diverse stomatal features across species, we incorporated additional datasets into model training. The 30 images in the maize dataset (Aono *et al*., 2021) represent a unique stomatal arrangement of monocots with subsidiary cells and narrow apertures. Additionally, glue imprints from *Zea mays* populations were included to capture variations in ploidy and cell morphology induced by colchicine treatment (Aono *et al*., 2021). We also included an in-house dataset of 31 images of *Lilium orientalis* cv. *Stargazer* to represent additional features of monocot stomata (Wu, 2016). Additionally, we extracted a supplementary dataset of 147 images from the LeafNet training set (Fig.1, Table S1) (Li *et al*., 2022). This further improved the ability of our model to adapt to a wide range of stomatal features and complex backgrounds.

### Few-shot transfer learning and average precision

YOLOv7-seg is the foundational model of StomaVision because of its robust capability for feature extraction and small-object identification(Wang *et al*., 2023a). However, the initial training of YOLOv7 was not based on the plant stomatal images. To address this gap, the model was fine-tuned using a dataset consisting of 337 stomatal images. Comparative evaluations with retrained Detectron2 (Wu *et al*., 2020) showed that the fine-tuned YOLOv7-seg outperformed its counterpart, achieving a 78.6% enhancement in bounding box average precision (bAP50) and 49.2% improvement in mean average precision (mAP50) (Fig. 2a,b, Table S2). Additionally, the predicted stomatal density was highly correlated with the manually labeled dataset (Pearson’s correlation coefficient test, *R* = 0.91, *P* < 2.2e^−16^) (Fig. 2c). Compared with inexperienced students labeling stomata images, StomaVision can produce more accurate results (Fig. 2d).

**Fig. 2.**
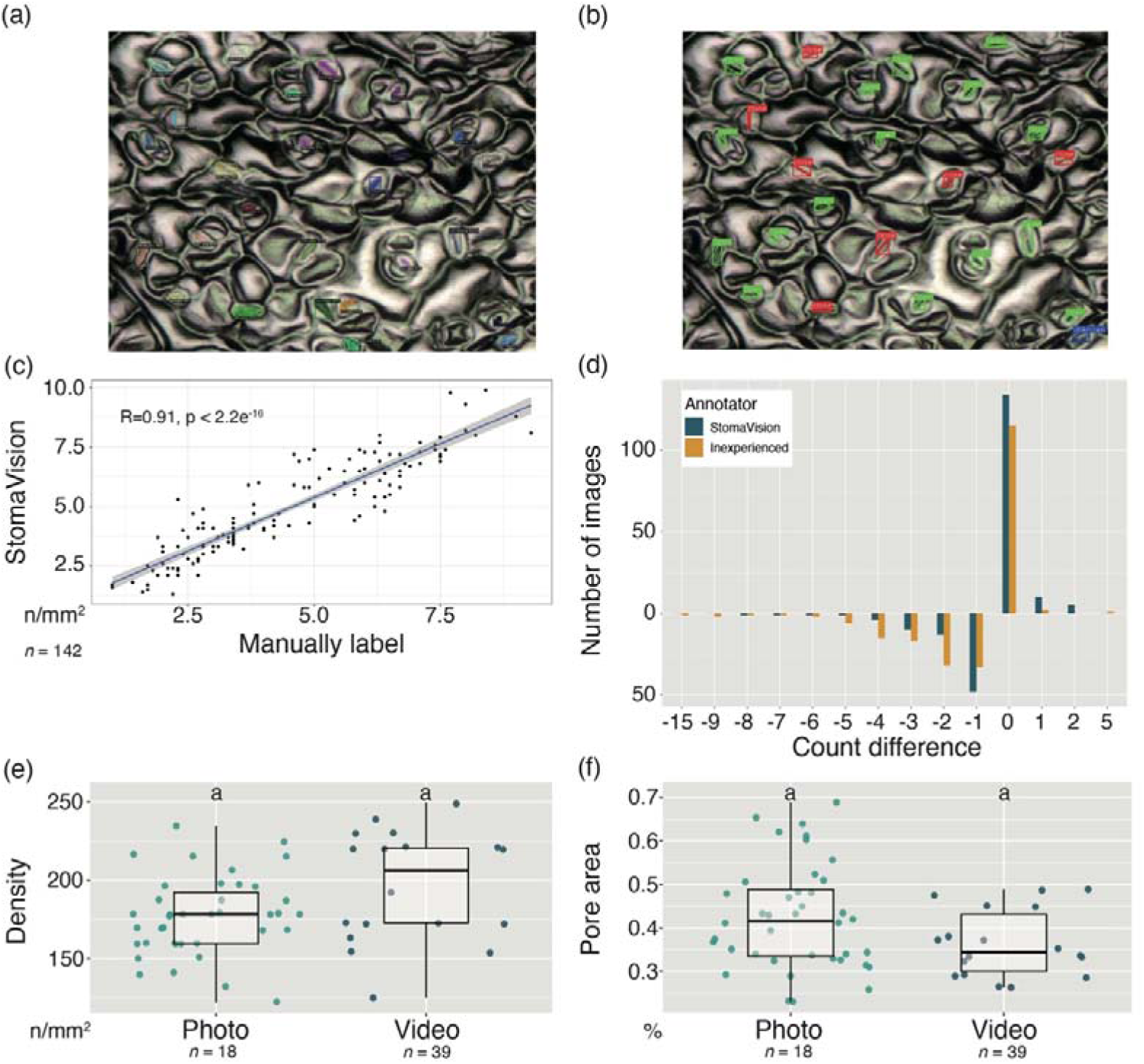
StomaVision prediction accuracy evaluation. (a) Stomata detection by StomaVision compared with (b) annotations by human experts. (c) Correlation analysis of stomatal density estimations between StomaVision predictions and expert evaluations of cauliflower samples. (d) Discrepancies in stomatal counts: comparison between StomaVision, human experts, and novices with basic stomatal knowledge, where negative values indicate undercounting, zero represents agreement, and positive values denote overcounting by StomaVision or novices. (e) Comparative analysis of ellipse and polygon methods for stomatal pore area measurement. (f) Regression analysis assessing the accuracy of the ellipse and polygon masking techniques in determining the pore area. (g-h) Comparative analysis of stomatal density and pore area measurements using two different input formats, photographs and videos, for the same stomatal micrographs.

Upon evaluation using the extensive ‘All’ dataset (Table 2), StomaVision exhibited a recall of 75.17%, precision of 80.80%, and F1-score of 77.88% (Table 3). The model effectively demonstrated its ability to accurately measure the stomatal aspect ratios using rotated bounding boxes. The precision of these measurements was assessed using the mean measurement errors, which generally remained below 0.2, reflecting a high level of accuracy in the computed dimensions. However, it is worth mentioning that the performance on the ‘Maize’ dataset was lower, with mean measurement errors exceeding 0.3. The high precision but low recall score of the ‘Maize’ dataset indicates that stomata detected by StomaVision are correct but likely missed stomata in the image (Table 3). The low recall of the ‘Maize’ dataset is mainly due to the small stomatal pores under low magnification rate of this dataset. The total stomatal pore area accounted for less than 0.5% of the total image area.

**Table 2.**
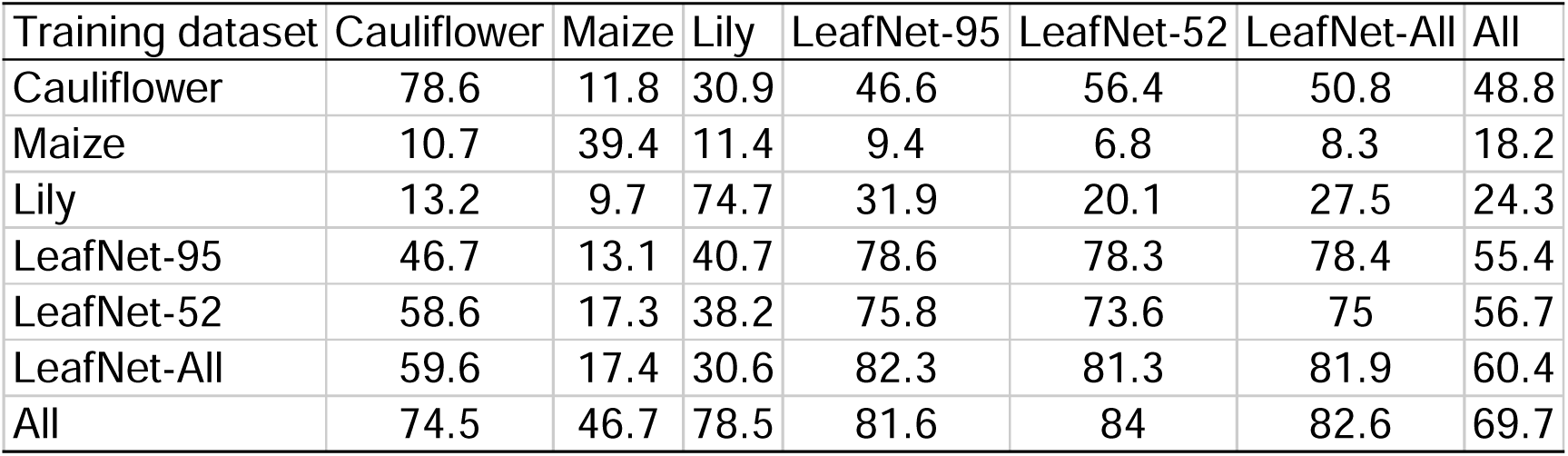
Few-shot performance of bounding box average precision 50 (bAP50) using StomaVision on different datasets.

| Training dataset | Cauliflower | Maize | Lily | LeafNet-95 | LeafNet-52 | LeafNet-All | All |
| --- | --- | --- | --- | --- | --- | --- | --- |
| Cauliflower | 78.6 | 11.8 | 30.9 | 46.6 | 56.4 | 50.8 | 48.8 |
| Maize | 10.7 | 39.4 | 11.4 | 9.4 | 6.8 | 8.3 | 18.2 |
| Lily | 13.2 | 9.7 | 74.7 | 31.9 | 20.1 | 27.5 | 24.3 |
| LeafNet-95 | 46.7 | 13.1 | 40.7 | 78.6 | 78.3 | 78.4 | 55.4 |
| LeafNet-52 | 58.6 | 17.3 | 38.2 | 75.8 | 73.6 | 75 | 56.7 |
| LeafNet-All | 59.6 | 17.4 | 30.6 | 82.3 | 81.3 | 81.9 | 60.4 |
| All | 74.5 | 46.7 | 78.5 | 81.6 | 84 | 82.6 | 69.7 |

**Table 3.** StomaVision performance on stomata detection.

| Dataset | Precision | Recall | F1-Score |
| --- | --- | --- | --- |
| Cauliflower | 0.7403 | 0.7396 | 0.7400 |
| Maize | 0.9451 | 0.2794 | 0.3946 |
| Lily | 0.8058 | 0.8293 | 0.8174 |
| LeafNet-All | 0.8400 | 0.8094 | 0.8244 |
| All | 0.8080 | 0.7517 | 0.7788 |

### Generalization capability and precision of StomaVision

The generalizability of machine learning algorithms is often limited by the diversity of training data. Given the heterogeneity of stomatal structures and variations in imaging techniques, the development of universally applicable machine-learning algorithms for stomatal image analysis is challenging. To test the generalization capability of StomaVision, we conducted evaluations that focused on its zero-shot learning capabilities. Zero-shot performance refers to the model’s ability to generalize to previously unseen data; in this case, stomatal images from species not included in the training dataset. Multiple independent training and validation datasets were generated by incorporating stomatal images from a range of species and imaging conditions. The zero-shot performance was measured in terms of the bounding box average precision (bbox AP50) and mask average precision (mask AP50) (Tables 3 and 4).

**Table 4.** Correlation of stomata anatomical traits with gas exchange analyzer.

|  | E | TleafCnd | VPDleaf | EBSum |
| --- | --- | --- | --- | --- |
| Pore axis ratio | NS | -0.41* | NS | NS |
| Stomata density | -0.51** | NS | NS | 0.36 * |
| Pore area/ total image area | -0.41* | 0.49** | 0.38 * | NS |
| Pore area | 0.38 * | NS | NS | -0.38 * |

Kendall rank correlation coefficient. NS not significant, \* $P < 0.1$ , \*\* $P < 0.05$

Models trained on comprehensive datasets, as well as those trained and validated on identical datasets, were considered soft upper bounds for performance (Tables 3 and 4), indicating the potential for improvement through more diverse and higher-quality training data. To enhance the dataset diversity, the LeafNet-All dataset was subdivided into two smaller datasets, namely LeafNet-95 and LeafNet-52, which contain 95 and 52 subset images, respectively. Both were designed to have validation sets that were subsets of the original LeafNet-All validation set. Overall, StomaVision exhibited strong zero-shot learning performance, particularly when fine-tuned using the LeafNet datasets (Tables 3 and 4). Through optimal hyperparameter tuning, the model fine-tuned on LeafNet-52 achieved the highest segmentation accuracy across all stomatal types, with bbox AP50 and mask AP50 scores of 84.0% and 53.7%, respectively. This was likely due to the inclusion of a diverse range of stomatal images from various species in the LeafNet dataset. However, it is worth noting that the generalizability of the model was less optimal when trained on datasets such as Maize and Lily, which presented challenges owing to the small size of stomatal pores in the input images.

Given the short inference time of the YOLOv7 architecture, StomaVision can complete a stomata prediction of 640 × 384 pixels in 10-15 ms per image. We further extended the inference to the video format and did not observe any additional overhead introduced by the video stream. The predicted stomata density (177±25.3 and 196±36.1 in photo and video, respective) and stomata pore area (0.420±0.116 and 0.364±0.077 in photo and video, respective) are compatible between input sources (Fig. 2e,f)

### Application of StomaVision in understanding heat stress responses

The robustness and generalization of StomaVision are illustrated by efficiently processing large datasets of stomatal images and video data without retraining with species-specific stoma images. This tool has facilitated quantification of stomatal traits and can be used in large-scale studies. Stomatal traits are modulated by the environmental conditions and global warming has seriously affected food security, therefore, we investigated how temperature influences stomatal physiology and plant growth. As stomatal features are influenced by both their genetic background and the growth environment, we compared these traits between genotypes. The genotype H37 (Ching Long Seed Co. Ltd., Taiwan) is suitable for cauliflower production in the tropical region at 26-35°C, featuring smaller plant architecture, a matuer curd size of 0.6 kg, and can be harvested in 37 days after transplantation. Coversely, the genbotype S100 (Ching Long Seed Co. Ltd., Taiwan) is suitable for the temperate region at 8-16°C, with large plant architecture, a matuer curd size of 2.3 kg, and can be harvested in 100 days after transplantation.

To reveal the interaction between genotypes and stomatal physiology in relation to the heat response, we treated H37 and S100 with either short-term (2-4 days, ST) or long-term (5-10 days, LT) high-temperature exposure (40°C). We then used StomaVision combining with the ‘gold-standard’ infrared gas exchange analyzers to assess the effects of different heat stress durations on cauliflower varieties by comparing stomatal traits. This approach overcame the traditional limitations associated with time-consuming micrograph analysis and the low throughput and labor-intensive use of infrared gas exchange analyzers for physiological measurements. Since stomata are gateways for water evaporation to reduce body temperature, we first evaluated the body temperatures of control and heat-treated plants. We observed an increase in leaf canopy temperatures in both H37 (23.5±1.54, 31.6±1.39 and 41.5±4.34°C respectively, Fig. 3) and S100 (23.2±1.28, 31.1±1.03 and 37.8±5.04°C respectively, Fig. 3) in ST and LT groups.

**Fig. 3.**
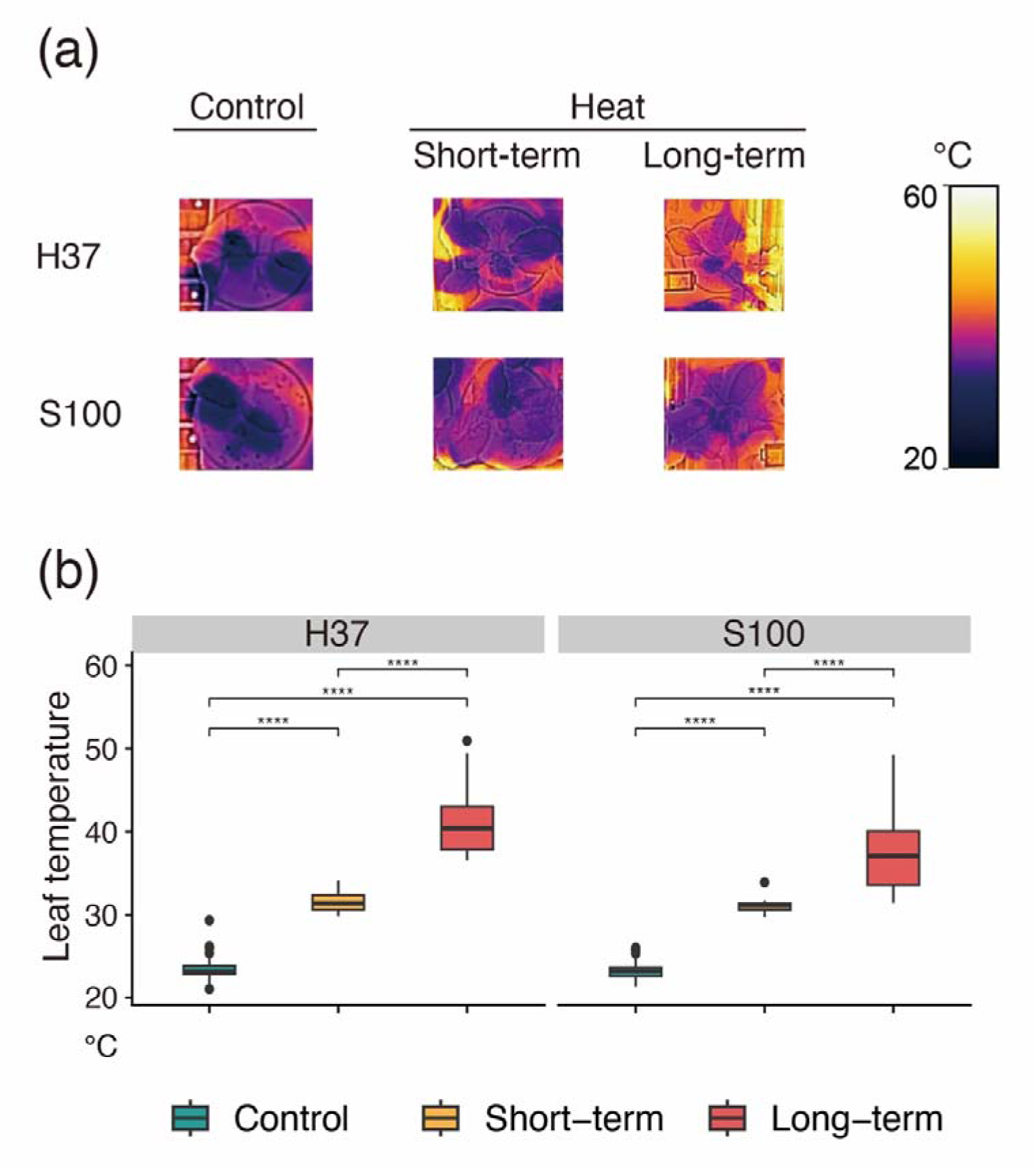
Leaf thermal temperature. (a) Leaf temperature in infrared thermal imagery. (b) Variations in the leaf temperature of genotypes H37 and S100 in the control and heat stress groups. The leaf temperatures of H37 and S100 were measured in 36, 12, and 18 plants in the control, ST, and LT groups, respectively. Horizontal lines within boxes indicate the median for each temperature based on measurements within a group. Boxes indicate the upper (75%) and lower (25%) quartiles. The dots represent outliers. Statistical significance was assessed using the Kruskal-Wallis test, where *** *P*<0.001.

Closing stomata is a rapid method employed by plants to minimize water loss. Nevertheless, plants adjust their developmental processes to withstand heat stress and ensure survival. To investigate how stomata respond to prolonged heat exposure, we analyzed parameters such as stomatal density, pore axis ratio, and pore area (Fig. 4a-c). Stomatal density was reduced in H37 under LT condition (230±91.9, 239±97.6, and 211±89.3 stomata per mm^2^ in the control, ST, and LT groups, respectively). But it was increased in S100 (222±98.9, 229±69.8, and 268±144.0 stomata per mm^2^ in the control, ST, and LT groups, respectively) (Fig. 4c). The result suggests that H37 and S100 may modulate stomatal development differently, and S100 does not as tightly as H37 to form stomata. Therefore, S100 had a broader distribution in stomatal density under LT treatment than H37 did (268±144.0 versus 211±89.3 stomata per mm^2^ in S100 and H37, respectively). The stomatal aperture axis ratio of H37 (0.390±0.124, 0.391±0.126 and 0.380±0.122 in control, ST and LT group, respectively) and S100 (0.428±0.133, 0.385±0.125 and 0.387±0.140 in control, ST and LT group, respectively) (Fig. 4d) were decreased under the long-term heat treatment. Pore area of H37 (1.22±0.441, 1.10±0.330 and 1.08±0.333 µm^2^ in control, ST and LT group, respectively) and S100 (0.933±0.380, 0.986±0.253 and 0.903±0.453 µm^2^ in control, ST and LT group, respectively) (Fig. 4e) also showed the smaller sizes in the LT group compared to the control. These data suggest that in response to long-term heat stress, plants mitigate water loss by reducing stomatal pore size, which increases leaf temperature (Fig. 3). The dynamic pattern of treated groups also demonstrates a constant adjustment of stomatal development and movement to cope with changing environmental conditions. Therefore, a high-throughput stomatal image analysis offers valuable information on genotype-specific adaptability to thermal stress.

**Fig. 4.**
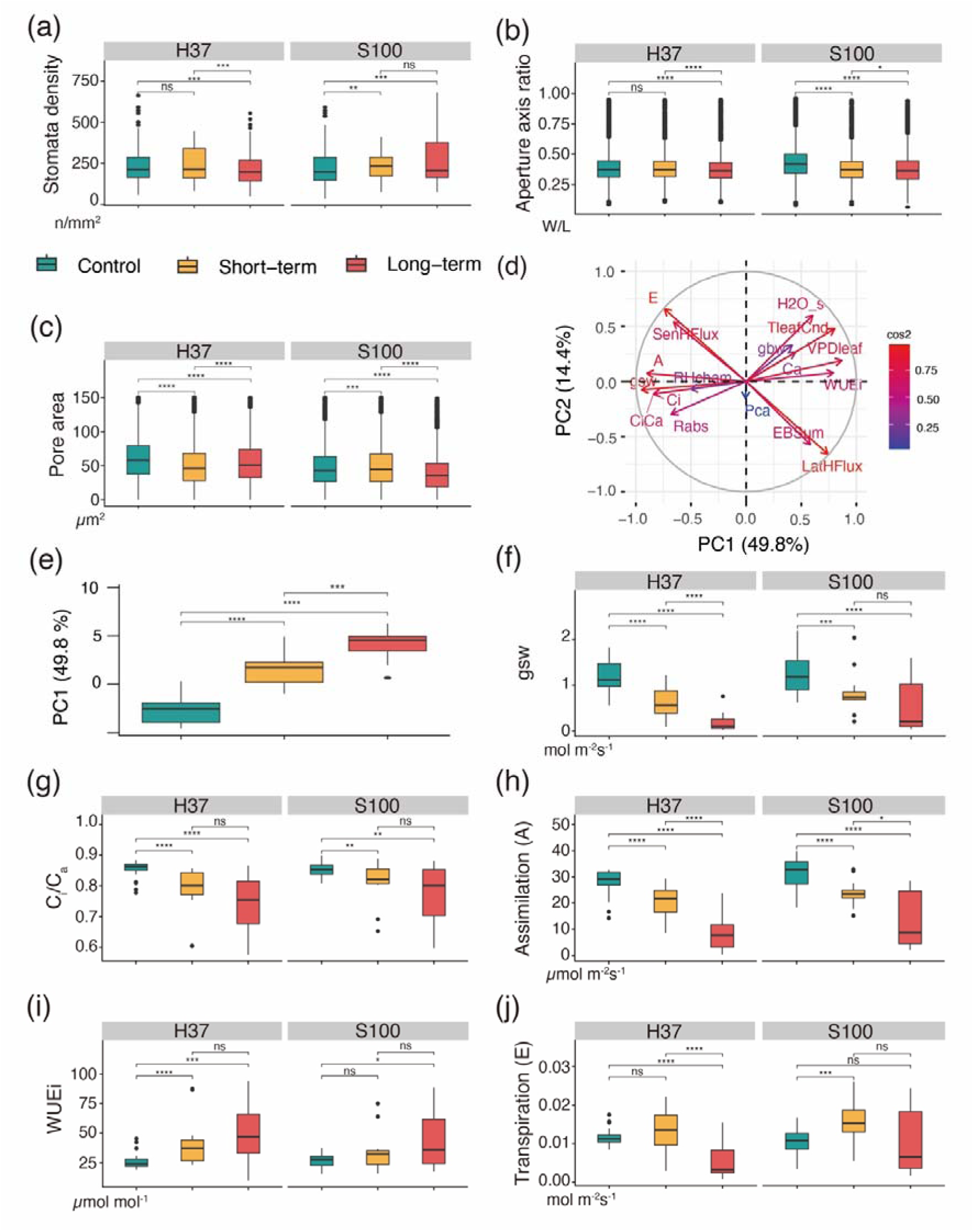
Plant trait analysis of genotypes H37 and S100 in the control and heat stress groups. Leaf micrographs of (a) stomatal density (H37 n=615, 280, and 396, and S100 n=640, 276, and 368 images in the control, ST, and LT groups, respectively), (b) stomatal aperture axis ratio (short/long axis) (H37 n=10,501, 4,324, and 5,694, and S100 n=8,887, 4,278, and 7,488 stomata), and (c) representative pore area per image (H37 n=10,501, 4,324, and 5,694, and S100 n=8,887, 4,278, and 7,488 stomata) were analyzed using StomaVision. (d) Principal component analysis (PCA) performed on the studied traits measured using LI-COR 6800 showed that the samples were significantly separated by photosynthesis and gas exchange. (e) The first PCA illustrates the differences between the control, ST, and LT groups. Boxplots of (f) stomatal conductance to water vapor (*gsw*), (g) leaf intercellular to atmospheric CO_2_ concentration ratio (Ci/Ca), (h) assimilation rate (A), (i) intrinsic water use efficiency (WUEi), and (j) transpiration rate (E) with n=35, 17, and 15 in H37, and n=45,13 and 17 in S100 in the control, ST, and LT groups, respectively. Horizontal lines within boxes indicate the median for each trait based on measurements within a group. Boxes indicate the upper (75%) and lower (25%) quartiles. The dots represent outliers. Statistical significance was assessed using the Kruskal-Wallis test, where *ns* was non-significant, *, *P*<0.05, **, *P*<0.01, and ***, *P*<0.001.

To further explore the contribution of parameters assessed within the gas exchange and photosynthesis system derived from the gas exchange analyzer, we conducted a principal component analysis and the correlation coefficient analysis based on the collected measurements in the control and treated groups (Tables 4 and S1). We identified that stomatal traits derived from the stomatal image analysis exhibited both strong positvie and negative correlcations with parameters assessed by the gas exchange analyzer (Table 4). We found a substantial variance primarily characterized by traits associated with photosynthesis, stomatal anatomy, and stomatal functionality (Xie *et al*., 2022), capturing 48.9% of the total variability (PC1 in Fig. 4d). The first axis (PC1) was mainly described by nine traits namely stomatal conductance to water vapor (gsw), leaf intercellular to atmospheric CO_2_ concentration ratio (Ci/Ca), assimilation rate (A), intrinsic water use efficiency (WUEi), transpiration rate (E), sample cell H_2_O concentration (H2O_s), leaf temperature for gas exchange (TleafCnd), vapor pressure deficit at leaf temperature (VPDleaf) and sum of energy balance components (EBSum) (Table S1), that showed high variability among the heat stress groups (Fig. 4e). Stomatal conductance to water vapor (*gsw*) was influenced by stomatal morphology, such as density and pore size. In line with our observations, the gsw in the ST and LT groups was reduced compared to that in the control group in H37 (1.180±0.333, 0.644±0.333, and 0.187±0.200 mol m^−2^s^−1^ in the control, ST, and LT groups, respectively) and S100 (1.250±0.436, 0.834±0.468, and 0.515±0.530 mol m^−2^s^−1^ in the control, ST, and LT groups, respectively) (Fig. 4f). The wide range of gsw distribution observed in the S100 LT groups could be attributed to the significant variance in stomatal density (Fig. 3a), given that stomatal density is a factor influencing *gsw*. However, despite this variability, both H37 and S100 groups exhibited a decreasing trend in gsw under heat treatments. In addition, we analyzed the net rate of CO2 assimilation (A) and intrinsic water use efficiency (WUEi) to understand the physiological outcomes of altering stomatal density and pore size. The internal to atmospheric CO_2_ concentration ratio (Ci/Ca) (H37 0.855±0.026, 0.784±0.075, and 0.747±0.089, and S100 0.853±0.0218, 0.811±0.0672, and 0.772±0.0930 in the control, ST, and LT groups, respectively) (Fig. 4g) and CO_2_ assimilation rate (A) (H37 28.0±4.67, 21.2±5.67 and 8.3±6.74 µmol m^−2^s^−1^, and S100 31.4±5.53, 23.8±5.14 and 14.1±10.50 µmol m^−2^s^−1^ in the control, ST, and LT groups, respectively) (Fig. 4h) were reduced by the heat treatment.

Although the Ci/Ca ratio was slightly reduced in the heat treatment groups (Fig. 4g), the carbon assimilation rate had a more pronounced decreasing (Fig. 4h), suggeting that in addition to the carbon concentration difference, the enzymatic activity of the photosynthesis system under heat treatment was much impaired. The intrinsic water use efficiency (WUEi), which is calculated by A divided by gsw (A/gsw) and reflects photosynthetic efficiency irrespective of environmental stimuli, demonstrated higher values in the treated groups compared to the control group (H37 26.5±6.23, 41.5±18.9, and 49.1±23.4 µmol m^−2^, and S100 26.8±5.78, 35.0±16.7, and 43.7±22.1 µmol m^−2^ in the control, ST, and LT groups, respectively) (Fig. 4i). This suggests that despite a decrease in carbon assimilation (Fig. 4h), the impact of stomatal conductance (gsw) was significantly enhanced under heat treatment. This enhancement could be attributed to factors such as a reduction in stomatal density, pore size, or alterations in stomatal dynamics induced by heat stress (Fig. 4a-c). The transpiration rate (E) of plants was observed to be reduced in the heat-treated plants to eliminate water loss while increasing body tempature (H37 0.0116±0.00198, 0.0134±0.00512, and 0.0053±0.00429 mol m^−2^ s^−1^, and S100 0.0102±0.0038, 0.0156±0.00528, and 0.0103±0.00816 mol m^−2^ s^−1^ in the control, ST, and LT groups, respectively) (Fig. 4j). Consistent with the observation of gsw (Fig. 4f), the decrease in E could be attributed to changes in stomatal traits, as plant transpiration is influenced by environmental temperature and surface characteristics like stomatal features.

The ultimate plant response showed a notable delay in curd induction time in the genotype H37 due to heat stress (27.3±0.98, 30.3±0.52, and 32.8±2.54 day in the control, ST, and LT groups, respectively) (*P* < 0.001) (Fig. 5a), underscoring the impact of vegetative stage stress on subsequent reproductive development. Since S100 is the winter genotype requiring vernalization, we could not observe its curd induction time here. The consequences of heat stress were evident in the varying growth rates of biomass. As two of the genotypes continued to be in the active vegetative growth phase during the heat experiment and H37 transitioned into the reproductive stage of curd induction, we analyzed the biomass at different growth stages (Fig. 5b,c). As anticipated, the LT group for H37 and S100 both exhibited significantly reduced biomass increase indices compared to the control group (H37 5.49±1.88 and 2.81±0.71, and S100 13.1±7.27 and 4.93±3.37 in the control, and LT groups, respectively) (Fig. 5d). The biomass increase index reflects the increase in biomass during the growth stage. Surprisingly, genotype H37 demonstrated a lower biomass increase index in the LT group than genotype S100. Additionally, genotype H37 exhibited a significantly greater biomass reduction index in the LT group, whereas the reduction in genotype S100 between the ST and LT groups was not significant (H37 −0.346±0.196 and −0.472±0.199, and S100 −0.432±0.191 and −0.424±0.182 in the ST and LT groups)(Fig. 5e). The biomass reduction index indicates biomass changes resulting from heat stress. Both H37 and S100 exhibited increased water use efficiency (WUEi, as shown in Fig. 4i) and decreased transpiration (E, as shown in Fig. 4j) under heat stress conditions. However, only H37 displayed greater biomass loss. Consequently, we hypothesized that despite the necessity for vernalization, S100 demonstrated superior responses and adaptation to prolonged heat stress.

**Fig. 5.**
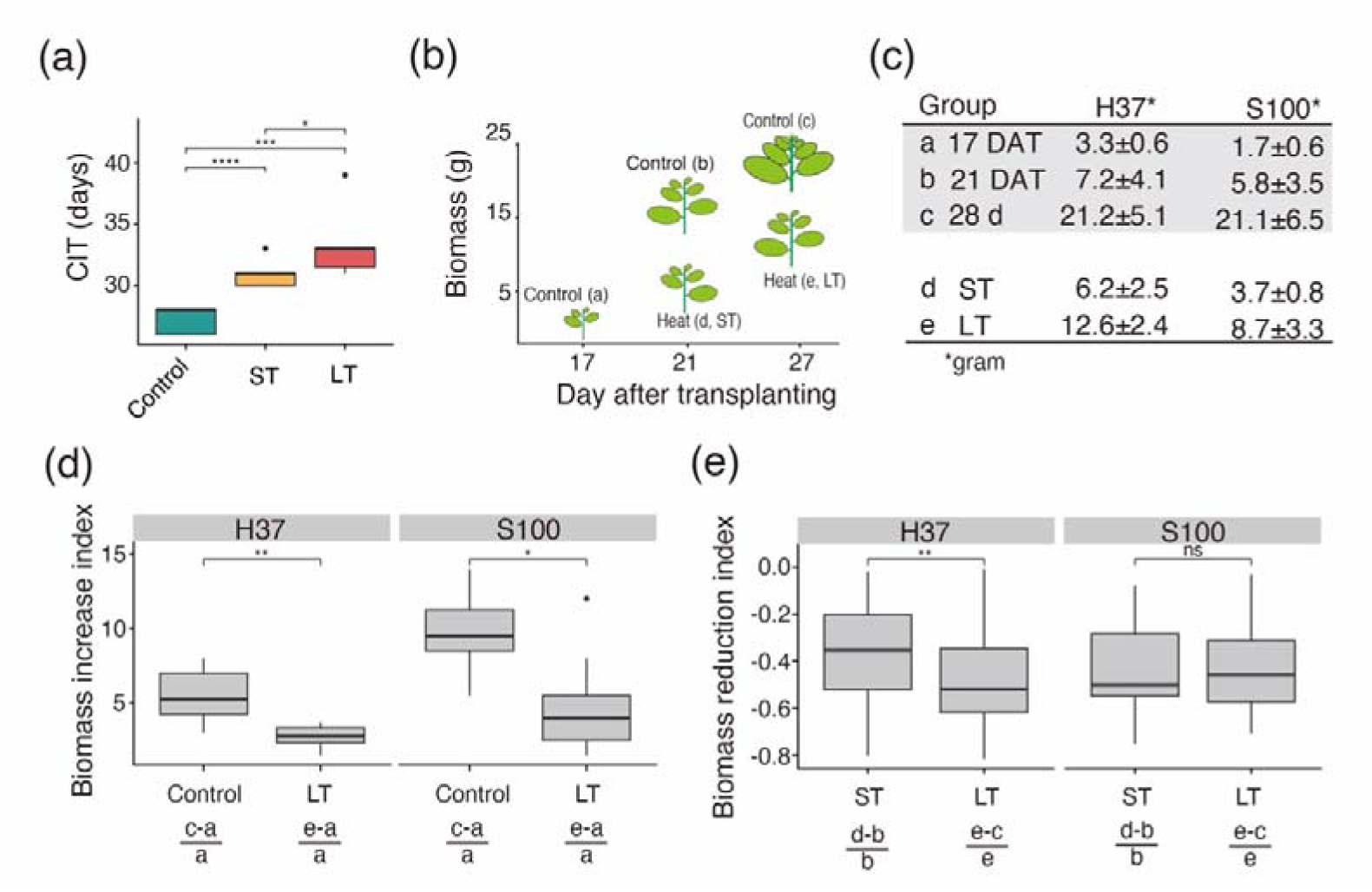
Phenotypic variations in genotypes H37 and S100 in the control and heat stress groups. (a) Curd induction time (CIT) was delayed in the ST and LT groups (n=33, 66, and 8 in the control, ST, and LT groups, respectively). (b) Illustration of different growth stages and heat stress groups. Plant age was calculated based on the days after transplanting (DAT) (c) Measured biomass during the different DAT and heat treatment groups. (d) Difference in biomass increase indices between the control and LT groups in the two genotypes (n=9 in each comparison). The biomass increase index reflects the increase in biomass during the 10 days of plant growth. (e) Difference in biomass reduction index between ST and LT groups in two genotypes (n_H37_=46, 53 and n_S100_=23, 30 in the ST and LT, respectively). The biomass reduction index indicates biomass changes resulting from heat stress. The more negative the value, the greater the damage caused by the heat stress. Statistical significance was assessed using the Kruskal-Wallis test, where *ns* was non-significant, *, *P*<0.05, **, *P*<0.01, and ***, *P*<0.001.

### Deployment of StomaVision in a container environment and web-based user interface

To facilitate broad accessibility and ensure adaptability to rapidly evolving machine learning frameworks, StomaVision was built on the foundation of the open-source Versatile Data Pipeline (VDP). A web-based user interface of StomaVision has been developed and is publicly available at https://stomavision.streamlit.app/ for users seeking expedited stomatal image analysis. For those requiring batch processing of large datasets, the complete environment is accessible via a command line interface. The system is containerized via Docker, allowing for both portability and ease of installation. Comprehensive setup instructions are available in the accompanying readme file, enabling users to establish a local Docker environment and web server (https://github.com/YaoChengLab/StomaVision). This local setup permits the batch processing of extensive image datasets via the command line interface, thereby offering a scalable solution.

StomaVision was designed to operate as a generalized model that is compatible with a wide array of stomatal images. This eliminates the need for users to specify image types or the species from which the images are derived, thus simplifying the user experience. The system produces two distinct output categories.

1. **Annotated Images:** These contain dual bounding boxes for each stomatal complex to delineate both the stomatal guard cells and stomatal pore. The pores were further annotated with polygonal shapes to mark their positions accurately.
2. **Statistical Summary:** A tabular output provides quantifiable metrics, including stomatal counts in the image. Each annotated stoma in the image corresponded to a unique numerical identifier. This was accompanied by measurements of the longest and shortest axes of the stomatal pore as well as the calculated pore area (Fig. 1d).

In summary, StomaVision not only offers an accessible and adaptable platform for stomatal image analysis, but also provides a range of output options to cater to both general and specialized research needs.

## Discussion

In this study, we introduced StomaVision, a fully automated, end-to-end program designed for the precise measurement of stomatal traits. The program was underpinned by the deep neural network of the instance-segmentation model YOLOv7 (Wang *et al*., 2023a). This foundational architecture was fine-tuned to enhance the accuracy of the stomatal detection and measurement. StomaVision has been trained and evaluated across a variety of species, each exhibiting distinct stomatal characteristics under different microscopic settings. Using the real-time image recognition capabilities of the YOLOv7 architecture, we demonstrated a novel approach that involves recording videos instead of capturing individual images during stomatal micrograph scouting (Fetter *et al*., 2019; Sakoda *et al*., 2019; Aono *et al*., 2021; Jayakody *et al*., 2021; Zhu *et al*., 2021; Liang *et al*., 2022; Pathoumthong *et al*., 2023; Sai *et al*., 2023). This approach substantially reduces the time required to obtain stomatal micrographs (Bourdais *et al*., 2019; Sai *et al*., 2023). In addition, StomaVision is equipped with a user-friendly web interface, and we provide comprehensive instructions for local training and installation, thereby ensuring sustainable accessibility and facilitating the ongoing development of the tool.

StomaVision also exhibited a good generalization capability in stomatal trait analysis without the need to retrain the machine learning models (Table 3). Recent advancements in deep learning algorithms for stomatal detection have led to a more general approach that can be applied to various plant species (Eisele *et al*., 2016; Aono *et al*., 2021). Various strategies have been developed to adapt to specific needs during model training. Image preprocessing steps such as downsampling, resizing the image to a specific resolution, image enhancement, noise reduction, processing, and transformation of the image between different color spaces are frequently used (Toda *et al*., 2018; Gibbs *et al*., 2021; Jayakody *et al*., 2021; Li *et al*., 2022). In addition to manually labeling stomatal apertures, Aono *et al*. manually selected and labeled a subset of noisy images, such as trichomes and epidermal cells, with various image resolutions to classify them as stomata and non-stomata for model training (Aono *et al*., 2021). Instead of training on complete micrograph images, some models have been specifically trained on a single stoma per image to improve the accuracy of stoma segmentation (Li *et al*., 2019). To further improve model accuracy, some tools use additional filtering steps after the prediction from deep learning models to remove stomata, such as a predefined axis ratio, extremely small size (< 5 pixels), or statistical filtering of abnormal stoma traits in the same image (Jayakody *et al*., 2021; Sai *et al*., 2023). Developing a generalized model, such as StomaVision, will broaden the use of stomatal analysis for plant biologists.

To ensure that StomaVision could perform automatic end-to-end stoma prediction from diverse microscope settings, we did not apply image preprocessing or post-processing steps during model training and prediction. This resulted in overall good performance when the stomatal density was approximately 200 per mm^2^, and these pores accounted for approximately 0.5% of the total image area in the dicot dataset (Fig. 2 and 3, Table 3). Nevertheless, we encountered constraints when performing model inference on stoma features of small dimensions that were unevenly distributed in the image (Table 3). We obtained suboptimal performance when applied to the maize and lily datasets, where the image contained too few stomatal image areas (0.05% of the total stomatal pore pixel/total image pixel) (Table 4). This intrinsic limitation of the Mask R-CNN and Detectron2 (Wu *et al*., 2020) models was also observed in SAI, where the prediction of barley images of larger stoma size performed better than *Arabidopsis* images (Sai *et al*., 2023).

We used a suite of classic object detection metrics, including precision, recall, and F1-score, along with the widely used Average Precision (AP) metric, to conduct a comprehensive evaluation of the performance of StomaVision. This approach aligns with several studies that predominantly focus on counting stomata and approximating their localization (Duarte *et al*., 2017; Toda *et al*., 2018; Bourdais *et al*., 2019; Fetter *et al*., 2019; Sakoda *et al*., 2019; Li *et al*., 2022). Accordingly, the model outputs that correspond to the ground truth are classified as true positives. Overall, StomaVision exhibited a good performance profile except for the maize dataset (Table 4). StomaVision has encountered challenges in achieving an optimal balance between precision and recall. The high precision score reflects the model’s efficacy in accurately identifying stomatal axes, yet it also missed several stomatal apertures, resulting in a lower recall score (Table 5). The limitations of the maize dataset are twofold: not only is it constrained by a small sample size (30 images) but it also features narrow apertures, small stoma sizes, and predominantly closed stomatal states, all of which restrict the performance of StomaVision. Implementing additional keypoint annotations, such as labeling the four points of the two axes and integrating keypoint detection with segmentation, will enhance the prediction accuracy.

Nevertheless, the broader issue of limited datasets continues to be a significant challenge in various research domains, particularly in the field of plant biology, where most training datasets do not contain sufficient plant data (Qi & Luo, 2019; Wolf *et al*., 2022).

Our innovative video recording approach, instead of capturing individual micrographs, will greatly accelerate image acquisition and address the challenge of insufficient image numbers. Typically, an experienced scientist spends 15-30 minutes to capture 5-10 images while scouting a micrograph to identify areas with clear stomatal images (Bourdais *et al*., 2019). In contrast, our video recording approach does not require automatic staging and focusing mechanisms exclusive to high-end microscopes. Manual adjustments of the stage and focus proved to be adequate for generating short videos that covered extensive areas of the slide. This method allows for rapid preparation of images for statistical analysis, with the data quality being comparable to that obtained from individual images. Moreover, the computational overhead introduced by processing individual frames from a video was negligible. To further enhance the utility and accuracy of this approach, additional techniques such as Fourier transform analysis or the Laplacian method can be used to remove out-of-focus images prior to stomatal analysis (Shivakumara *et al*., 2011).

One major drawback of YOLOv7 is its training time, which is approximately twice that of Detectron 2. This is mainly owing to the sophisticated data augmentation techniques used by YOLOv7, which doubled the number of training samples compared with Detectron 2. Unlike basic augmentations, such as flips and rotations, YOLOv7 uses more complex techniques, such as mosaic, mixup, and post-mosaic affine transformations (Wang *et al*., 2023a), based on data augmentation profiles developed by Ultralytics (https://github.com/ultralytics/ultralytics). We selected the medium augmentation profile ‘YOLOv7-med’ as the baseline model for StomaVision and applied fine-tuning to adapt the pre-trained model for stomata detection and pore measurements. Incorporating YOLOv7 into StomaVision significantly improves the ability of the model to accurately detect and measure stomata, even in instances where these features are small and sparsely distributed throughout the images.

StomaVision primarily serves as Software as a Service (SaaS) through a web-based interface designed to offer an accessible user experience without the complexities of software installation and parameter configuration. It is tailored to ensure seamless interaction for users, eliminating the need to differentiate between dicot and monocot settings, specify expected stomatal sizes, or select types of microscope images. (Fetter *et al*., 2019; Li *et al*., 2022; Pathoumthong *et al*., 2023). One of the challenges in maintaining software is its dependence on external software packages, which can cause malfunctions when these dependencies are updated (Barker *et al*., 2022). To address this issue, StomaVision is packaged in a container environment to enhance its interoperability and reusability. Advanced users are encouraged to consult the installation instructions on GitHub to configure their local Docker instances for an optimized experience (https://github.com/YaoChengLab/StomaVision).

It is possible to incorporate a human-in-the-loop or interactive machine learning model to adapt to the specific needs of stomatal prediction (Berg *et al*., 2019; Pachitariu & Stringer, 2022; Wang *et al*., 2023b). Nevertheless, there is generally a tradeoff between training on a specialized dataset to achieve good performance or a generalized model without the need to retrain the model.

### Stomatal traits as indicators of plant response to heat stress

Stomata are important for regulating water loss in exchange for CO_2_ uptake and controlling plant body temperature to maintain optimal physiological performance (Franks & Casson, 2014). Our heat stress experiments revealed the intricate dynamics among stomatal traits, physiological conditions, genetic diversity, and varying durations of heat stress.

The analysis classified the cauliflower plants into three phenotypic response groups based on heat exposure. Under heat conditions, the plants displayed elevated canopy temperatures (Fig. 3) and delayed curd initiation compared to those in the control group, which is a key reproductive trait (Fig. 5c). In this study, heat stress in the vegetative phase delayed reproduction in the summer genotype H37, contradicting earlier reports that elevated temperatures adversely affect photosynthetic performance and plant growth (Dikšaitytė *et al*., 2019; Charng & Chen, 2023). Prolonged exposure to 40°C is likely detrimental, surpassing optimal growth conditions and delaying plant development (Persaud *et al*., 2022; Liu *et al*., 2023).

The use of the binary classification of stomatal open or closed states and the stomatal pore axis ratio as a metric can be insufficient for a comprehensive understanding of stomatal dynamics and their role in gas exchange and transpiration (Xie *et al*., 2022). Stomata can be very leaky and probably incompletely closed (Duursma *et al*., 2019). To address this, our study integrated measurements of stomatal pore area, offering a more subtle view when analyzed alongside the pore axis ratio. This approach is consistent with previous research indicating a negative correlation between stomatal size, density, and conductance variability (Franks *et al*., 2009; Fanourakis *et al*., 2015). By including additional parameters from the gas exchange analysis, our results support models that correlate stomatal conductance with photosynthesis (Franks *et al*., 2009; Damour *et al*., 2010). Our findings revealed that under heat stress, plants reduced stomatal numbers and pore size, leading to decreased transpiration and assimilation rates. Altered stomatal conductance (*g_sw_*), and decreased intracellular to ambient CO_2_ concentrations (Ci/Ca), ultimately enhanced water use efficiency (WUEi) by minimizing water loss.

Previous studies in *Arabidopsis* have shown that the bHLH transcription factor PHYTOCHROME-INTERACTING FACTOR 4 (PIF4) is activated at high temperatures, which binds and represses the expression of *SPEECHLESS (SPCH)* and *MUTE*, and lowers stomatal differentiation rates (Lau *et al*., 2018; Endo & Torii, 2019; Samakovli *et al*., 2020b). Stomatal density in *Arabidopsis* under heat stress is modulated by the interaction of HEAT SHOCK PROTEINS 90 (HSP90s) and YODA mitogen-activated protein kinase kinase kinase, which affects the phosphorylation and deactivation of key transcription factors SPCH (Samakovli *et al*., 2020b). Unlike the decrease in stomatal density observed in H37 under heat treatment, S100 displayed higher stomatal density. This suggests that stomatal formation in the winter genotype S100 is less affected by the PIF4 regulatory pathway, as observed in H37 and *Arabidopsis*. Further investigations will be necessary to elucidate the underlying mechanism. Interestingly, our heat treatment of cauliflower shared similar patterns with *Arabidopsis* where different heat stress duration caused different stomatal density and stomatal index (Samakovli *et al*., 2020a). Our results demonstrated that with prolonged heat stress, *g_sw_* must be reset in new leaves by adjusting the number and size of the stomata. The heat stress duration likely caused different levels of impact on stomatal and physiological traits and highlighted the compensatory capacity of plants to cope with the stress environment (Almeida *et al*., 2023).

Comparative analysis of genotypes H37 and S100 revealed unexpected stomatal behavior and physiological responses under heat stress conditions. Contrary to the general assumption that genotype H37, typically cultivated in tropical regions with temperatures ranging from 26-35°C, possesses superior stomatal and physiological adaptations for high-temperature tolerance, our findings indicate otherwise. The growth of H37 was more adversely affected by heat stress compared to S100 (Fig. 4 and 5). Alterations of stomatal traits, gas exchange and photosynthesis system led to a more significant biomass reduction in H37 than in S100 under heat stress conditions. This discrepancy highlights the complexity of stomatal and physiological responses to heat stress and underscores the need for further research to unravel the underlying mechanisms in crop species.

## Conclusion

The development of this tool will democratize stomatal analysis, enabling a more extensive range of biologists to engage with this vital aspect of plant physiology and also enhance the precision and scope of the measurements obtained. This progression is essential for advancing our understanding of stomatal behavior, particularly in an era in which plant resilience and adaptation are of paramount concern.

## Data Availability

The source code, trained model, user installation and training guideline, and all the labeled images of leaf stomata are available at https://github.com/YaoChengLab/StomaVision. The web portal of extracting stomatal traits is available at https://stomavision.streamlit.app/.

## Author Contributions

TLW, PYC, XD, PLC, and YCL conceived the research. TLW, JYO, PXZ, YLW, RHW, TCH, CYL, and YCL conducted the field and growth chamber experiments. TLW, JYO, PXZ, YLW, and RHW produced cauliflower images and labeled all model training data annotations. PYC, XD, PLC, and WYL developed and implemented machine learning strategies and developed an inference model. TLW, PYC, XD, HW, and YCLset up the online demo website and wrote an online user guide. TLW, PYC, and YCL drafted the manuscript. TLW, PYC, CMH, and YCL edited and revised the final manuscript. All authors have read and approved the final manuscript.

## Acknowledgements

The authors thanks Yi-Mei Huang and Ya-Sui Huang for field works and technical assistance, summer intern students in the YCL lab for technical assistance, and Miranda Loney for English editing. This research was supported by the Innovative Translational Agricultural Research Program (AS-KPQ-107-ITAR-10, AS-KPQ-108-ITAR-10, AS-KPQ-109-ITAR-10, AS-KPQ-110-ITAR-03, AS-ITAR-111-L11102, AS-KPQ-111-ITAR-11107, AS-KPQ-111-ITAR-11207, and AS-ITAR-111-L11202) and Academia Sinica Institutional Fund to Y-CL.

## Supplementary Information

**Table S1.**
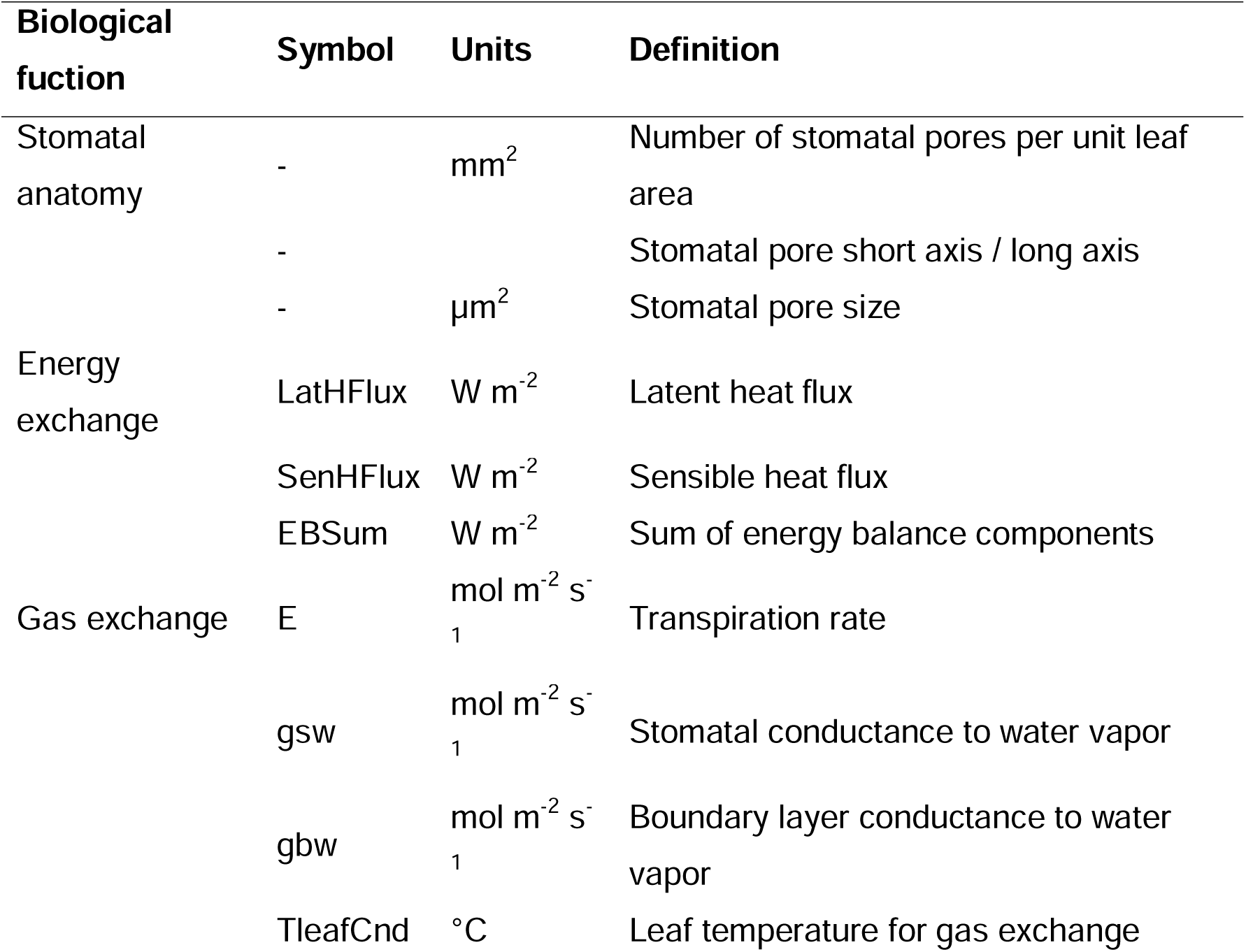
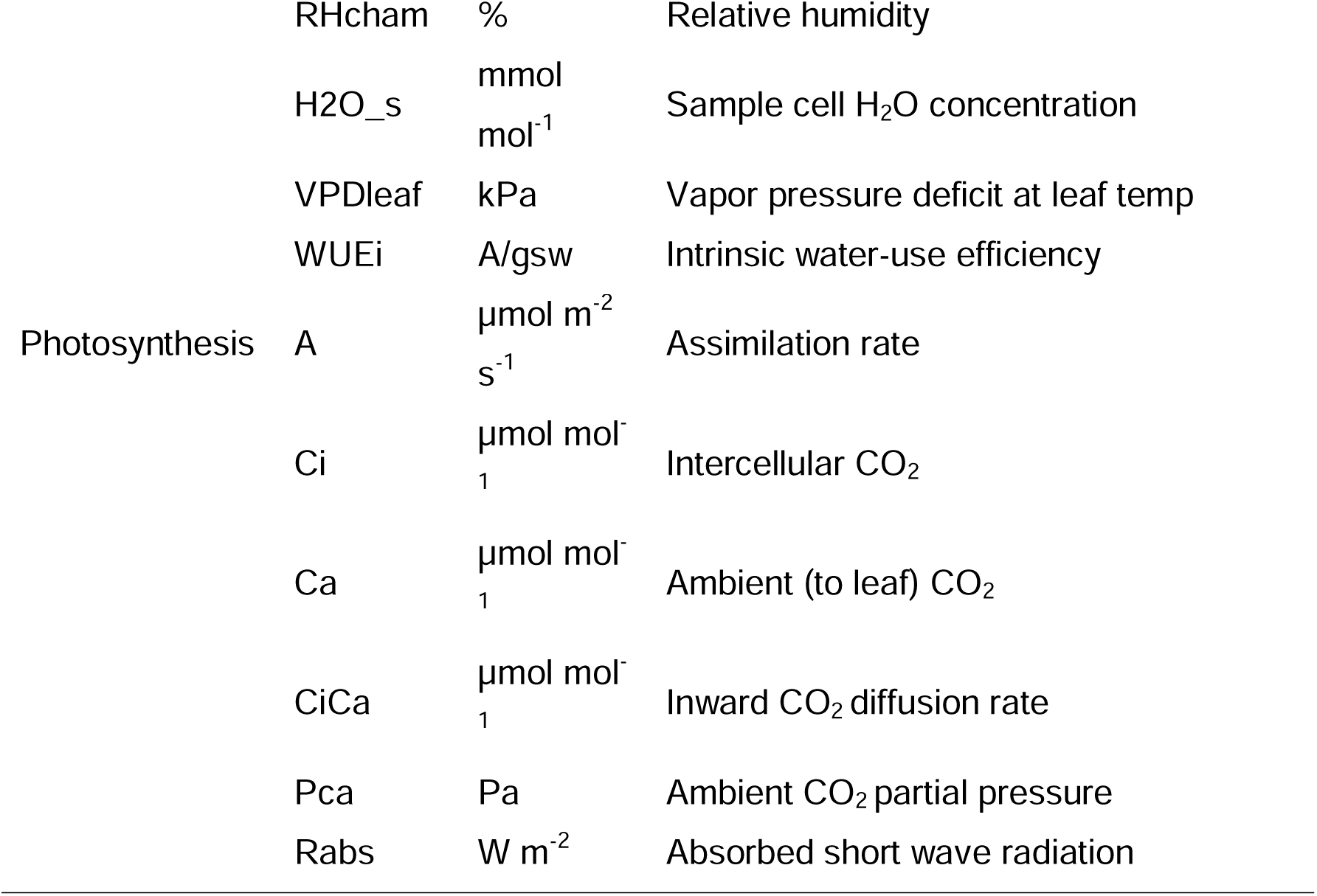
Difinition of traits and units used.

**Table S2.** Transfer learning performance with different base models.

| Base Model | Dataset | bAP(%) | bAP50(%) | mAP(%) | mAP50(%) |
| --- | --- | --- | --- | --- | --- |
| Detectron2 | Cauliflower | 29.2 | 64.2 | 13.9 | 38.0 |
| YOLOv7 | Cauliflower | 35.7 | 78.6 | 19.5 | 49.2 |

**Table S3.**
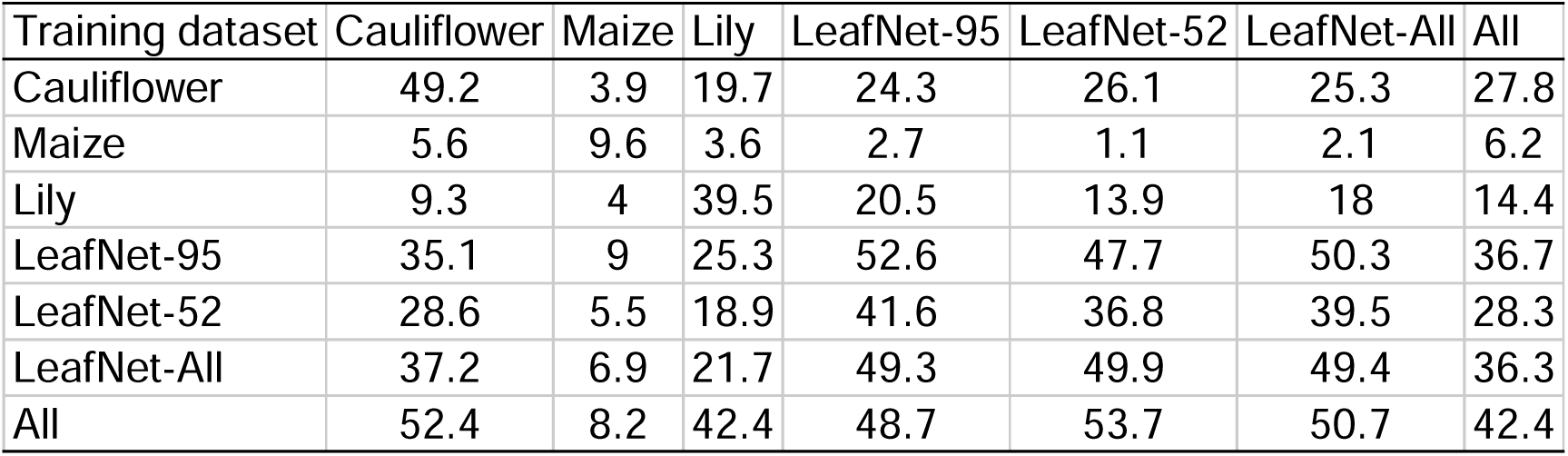
Few-shot performance of mask AP50 by StomaVision on different datasets.

**Fig. S1.**
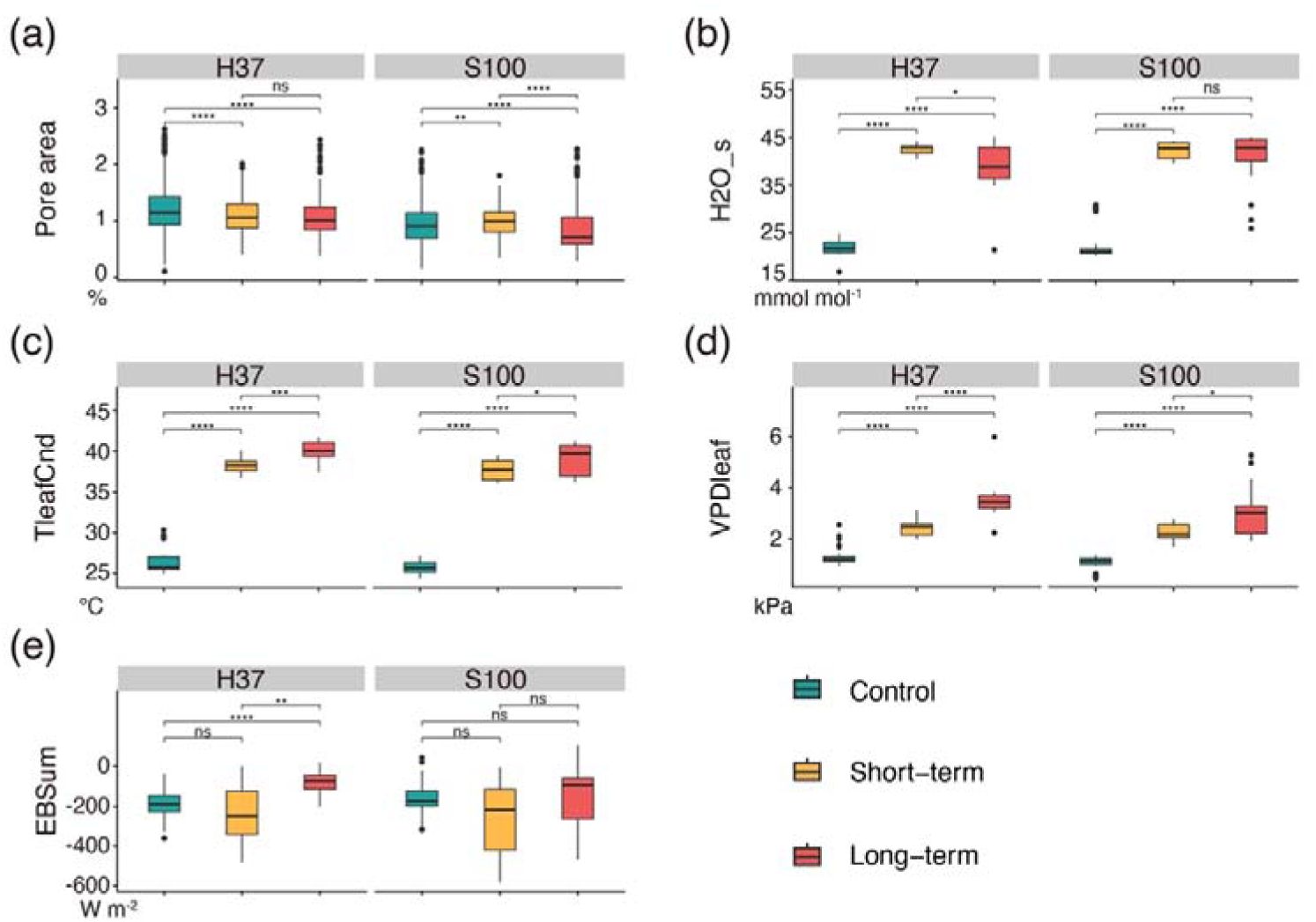
Variation in Physiological Traits. Boxplots of (a) sample cell H2O concentration. (b) Leaf temperature for gas exchange. (c) Vapor pressure deficit at leaf temperature and (d) sum of energy balance components in the control, ST, and LT groups (n_H37_=32, 17, and 15; n_S100_=45, 13, and 17). Horizontal lines within boxes indicate the median for each trait based on measurements within a group. Boxes indicate the upper (75%) and lower (25%) quartiles. The dots represent outliers. Statistical significance was assessed using the Kruskal-Wallis test, where *ns* was non-significant, *, *P*<0.05, **, *P*<0.01, and ***, *P*<0.001.

